# Altitudinal range-size distribution of breeding birds and environmental factors for the determination of species richness: An empirical test of altitudinal Rapoport’s rule and rescue effect on a local scale

**DOI:** 10.1101/398727

**Authors:** Jin-Yong Kim, Changwan Seo, Seungbum Hong, Sanghoon Lee, Soo Hyung Eo

## Abstract

Range-size distributions are important for understanding species richness patterns and led to the development of the controversial Rapoport’s rule and Rapoport-rescue effect. This study aimed to understand the relationship between species richness and range-size distribution in relation to environmental factors. The present study tested the following: (1) altitudinal Rapoport’s rule, (2) climatic and ambient energy hypotheses, (3) non-directional rescue effect, and (4) effect of environmental factors on range-size group. Altitudinal species range-size distribution increased with increasing altitude and showed a negative relationship with climatic variables and habitat heterogeneity, and a positive relationship with primary productivity. These results support the altitudinal Rapoport’s rule and climatic hypothesis; however, they do not fully support the ambient energy hypothesis. Results from testing the non-directional rescue effect showed that the inflow intensity of species from both directions (high and low elevations) affected species richness. And we found that the 2nd and 3rd quartile species distribution were the main cause of a mid-peak of species richness and the non-directional rescue effect. Additionally, the 2nd quartile species richness was highly related to minimum temperature and possessed thermal specialist species features, and the 3rd quartile species richness was highly related to habitat heterogeneity and primary productivity. Although altitudinal range-size distribution results were similar to the altitudinal Rapoport’s rule, the mid-peak pattern of species richness could not be explained by the Rapoport’s-rescue effect; however, the non-directional rescue effect could explain a mid-peak pattern of species richness.

## Introduction

To identify species richness patterns, analyses of geographical patterns in species richness have been traditionally performed [1,2]. However, a few decades ago, great attention was given to the necessity of studying species range-size distributions [2]. This concern regarding range-size distributions led to the development of the controversial Rapoport’s rule [3]. Rapoport’s rule states that higher latitudinal species have wider latitudinal ranges than that of lower latitudinal species and was developed by Stevens [3]. The effect of this phenomenon on species richness is explained by the rescue effect that suggests there is a source-sink dynamic in populations based on the influence of immigration on extinction [4], which is called the Rapoport-rescue hypothesis [3]. This phenomenon has been extended to an altitudinal gradient [5,6]. However, ever since this phenomenon was defined as a rule, there have been many associated controversies related to different results obtained for different taxa, sampling effort, geographical scale, and mechanism used [7–10].

One of the main underlying mechanisms of Rapoport’s rule is that species range-size distributions are determined by climatic conditions [3,6]. Organisms living at high latitudes or altitudinal areas, where climatic conditions are highly variable, have broader physiological thermal tolerances [11]. Therefore, this hypothesis proposes that organisms living in these areas will have a wider distribution range. Other ecological determinants of range-size distribution are associated with topographical complexity and habitat heterogeneity [12,13], which can be described by the ambient energy hypothesis. This hypothesis states that there exists a fine subdivision regarding limited food sources by the topographical habitat structure that promotes greater specialist species [14,15]. Thus, more frequent interaction among species could occur at higher latitudes, thereby species range-size distributions are wider with increasing latitude [15]. Although the importance of habitat heterogeneity is constantly mentioned together with climatic conditions, testing of the ambient energy hypothesis has not frequently occurred and there are many cases where habitat and topographical heterogeneity have not been distinguished from each other [13]. In addition, the altitudinal approach for this hypothesis has not yet been applied in advanced studies.

The predicted consequence of the altitudinal Rapoport’s rule is an increase in species richness from higher to lower elevations, which is termed the Rapoport-rescue hypothesis [6]. According to a study by Stevens [6], this phenomenon occurred in cases that showed decreasing patterns of species richness with increasing altitudes. The results from this study showed that low altitude localities had relatively more species near the edge of their range than high altitude sites [6]. Based on the hypothesis by Stevens, wide-range species are the main contributors to geographical patterns in species richness. However, Almeida-Neto et al. [16] stated the directions of three putative rescue effects in altitudinal gradients. Based on these, two hypotheses were formed as follows: when species richness exhibits a decreasing pattern with increasing altitude, an influx of species occurs at high or low altitudes, and when species richness indicates a mid-peak, the influx of species occurs via a non-directional rescue effect [16]. In the latter case, the increase in species richness might have occurred because of the other range species rather than because of the wide-range species. Therefore, different recue hypothesis should be applied based on different species richness patterns along altitudinal gradients. Thus, it is important to identify which range species increased the species richness and identify what environmental factors affect the distribution of each range species.

The present study aimed to understand the range-size distribution patterns and underlying mechanism along altitudinal gradients, and the relationship between species richness and range-size in relation to environmental factors. To understand the range-size distribution patterns and underlying mechanism along altitudinal gradients, we tested the altitudinal Rapoport’s rule and climatic and ambient energy hypotheses. Meanwhile, in a previous study, a hump-shaped pattern of species richness along an altitudinal gradient in the study site was identified [17]. Therefore, to determine the reason for such mid-peak patterns in species richness, we also tested the non-directional rescue effect and identified environmental factors associated with the range-size distribution group.

## Materials and methods

### Study site

The present study was conducted in mixed or deciduous forested areas in Jirisan National Park (South Korea), with altitudinal range from 200 to 1,400 m above sea level (asl). The altitudinal range in the park was from 110 to 1,915 m asl; however, we excluded the subalpine forest (up to 1,400 m asl) from the survey area to minimize the differences in bird communities among forest types [17].

### Species range-size distribution (Altitudinal Rapoport’s rule)

The distributions of 53 breeding bird species surveyed along an altitudinal gradient from a total of 142 plots were used from a database by Kim et al. [17]. To estimate the range-size distribution of each species, we identified the maximum and minimum altitude of each bird species distribution in a 100 m elevation band. Species that only occurred in a single plot were given a range of 100 m and included in the analysis. Then, the mean altitudinal range of species in a given plot was calculated by averaging the altitudinal range of each species present [6]. We identified the patterns in mean altitudinal range-size distributions using the best-fit curve (linear, quadratic, and exponential) estimation function in SPSS 20.

### Climatic and ambient energy hypotheses

To test the climatic hypothesis, the maximum and minimum temperatures during the 2015 breeding season were extracted from each survey plot using the Weather Research and Forecasting software program, version 3.6 [17].

Based on the ambient energy hypothesis, range-size distribution is related to primary productivity and habitat complexity [12,13]. Therefore, to test the ambient energy hypothesis, we used the vertical coverage of vegetation and horizontal habitat diversity as an indicator of primary productivity. The vertical coverage of vegetation classified the vertical layers into understory (< 2 m), midstory (2–10 m), and overstory (> 10 m) vegetation and included four categories in each layer : 0 (0% coverage), 1 (1%–33% coverage), 2 (34%–66% coverage), and 3 (67%–100% coverage) [17]. The horizontal habitat diversity was calculated by the Shannon-Wiener diversity index (H′) using the area of that particular habitat type (abundance) and the number of different habitat types (richness) [17], which was used as an indicator of habitat heterogeneity.

Analysis was conducted by model selection and multimodel inference using a generalized linear model. Before adding variables to the model, we identified the correlation between variables and eliminated maximum temperature correlated (r ≥ |0.7|) with minimum temperature (r = 0.991; S1 Table). We developed a set of seven candidate models and calculated Akaike’s information criterion adjusted for small sample sizes (AICc) and Akaike weights (*w*_*i*_) [18]. The high-confidence set of candidate models consisted of models with Akaike weights within 10% of the highest, which were used to compute model-averaged parameter estimates [17,19–21]. All statistical analyses were performed using the R Studio 1.1.383 software program (packages bbmle, AICcmodavg, and MuMin).

### Non-directional rescue effect (midpoint method)

To demonstrate the non-directional rescue effect, the midpoint of each species was calculated using the median of each species range [22,23]. Then, the distance between the two groups was calculated, which was divided by the center altitude (800 m asl) that showed the highest species richness [17] and averaged the midpoints of all the species detected at each group by the left and right group (> 800 m asl, < 800 m asl). In the case of a plot where species only existed on the left or right side, it was assumed that there was no influence of non-directional rescue effect and these species were given a value of 0. Using this method, we identified the intensity of species inflow along the distance from the center altitude. The relationship between the number of species and distance between the mean mid-point was analyzed using the best-fit curve (linear, quadratic, and exponential) estimation function in SPSS 20.

### Effect of environmental factors on the range-size distribution group (quartile method)

All bird species were divided using the quartile method based on their identified range-size distribution, i.e., less than 25% species (1st quartile species), between 25% and median number of species (2nd quartile species), between median number and 75% species (3rd quartile species), and more than 75% species (4th quartile species) [2]. To identify which quartile group increased the species richness, present/absent data of each quartile group were used and analyzed using the independent samples t-test in SPSS 20. The effects of environmental factors (climate, primary productivity, and habitat heterogeneity) were analyzed in the quartile groups that were determined to be affecting species richness. Analyses were conducted using model selection and multimodel inference in the R Studio 1.1.383 software program (packages bbmle, AICcmodavg, and MuMin) and the number of species for each quartile species.

## Results

### Species range-size distribution (Altitudinal Rapoport’s rule)

We tested the altitudinal Rapoport’s rule in 53 breeding bird species from 142 plots. The patterns in mean altitudinal range-size distributions showed a tendency of broader range-size distribution with increasing elevation (Fig 1). All curves (linear, quadratic, and exponential) represented by a significant relationship (*P* < 0.001, *P* < 0.001, *P* < 0.001, respectively; Fig 1). The highest value of R^2^ was a quadratic curve (R^2^ = 0.41; Fig 1.b); however, linear and quadratic curves showed a slight difference (Fig 1.a and b). The lowest value of R^2^ was an exponential curve (R^2^ = 0.38; Fig 1.c).

**Fig 1.**
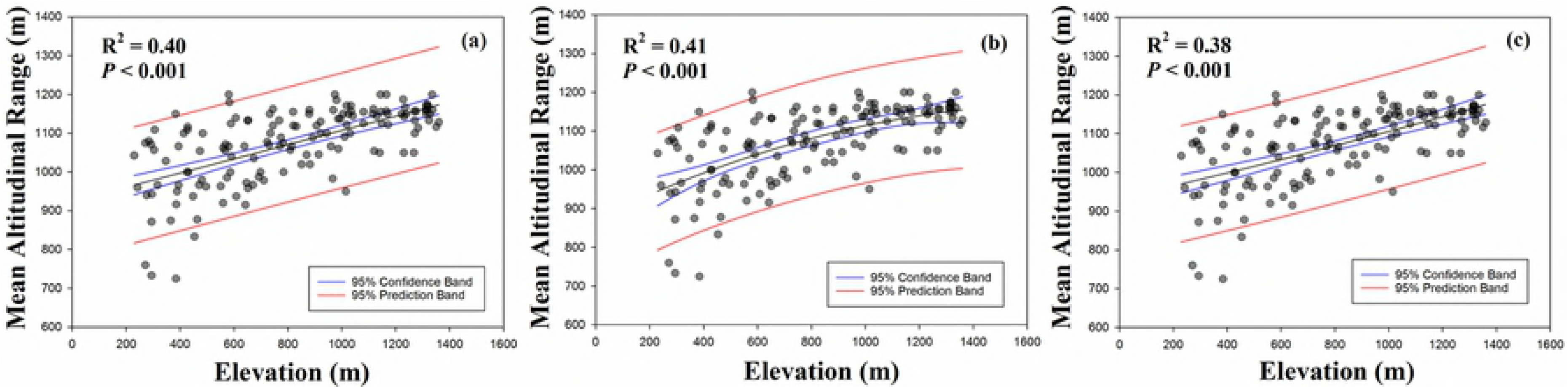
Altitudinal pattern of mean altitudinal range (m) in each plot. **(a)** linear, (b) quadratic, and (c) exponential curves were represented.

### Climatic and ambient energy hypotheses

To understand the underlying mechanism of Rapoport’s rule, we identified the influence of minimum temperature, vertical coverage of vegetation, and horizontal habitat diversity. The results from the model selection showed a set of candidate models with seven combinations of five variables showing two supported models that had Akaike weights within 10% of the highest weight (Table 1). The best model of mean altitudinal range included minimum temperature and habitat diversity (*w*_i_ = 0.870; Table 1). The second ranked model was the full model, which contained the added variable of vertical coverage of vegetation, in which the Akaike weights were 6.7 times lower than that of the best model (*w*_i_ = 0.870 vs *w*_i_ = 0.130; Table 1).

**Table 1.** Model selection for predicting mean altitudinal range based on minimum temperature, vertical coverage of vegetation, and habitat diversity. Coverage of understory vegetation = UV, midstory vegetation = MV, overstory vegetation = OV, minimum temperature = MT, habitat diversity = HD

| **Response variables** | **Candidate models** | **AICc** | **ΔAICc** | **df** | **$w_i$** |
| --- | --- | --- | --- | --- | --- |
| Mean Altitude Range | (Best model) Intercept + MT + HD | 1620.8 | 0.0 | 4 | 0.870 |
|  | (Full model) Intercept + MT + OV + MV + UV + HD | 1624.6 | 3.8 | 7 | 0.130 |
|  | Intercept + MT | 1635.2 | 14.4 | 3 | <0.001 |
|  | Intercept + MT + OV + MV + UV | 1637.3 | 16.5 | 6 | <0.001 |
|  | Intercept + OV + MV + UV + HD | 1639.8 | 19.0 | 6 | <0.001 |
|  | Intercept + HD | 1640.1 | 19.3 | 3 | <0.001 |
|  | Intercept + OV + MV + UV | 1694.4 | 73.7 | 5 | <0.001 |

Multimodel averaged parameter estimates including the two supported models over the mean altitudinal range showed negative correlation with minimum temperature and habitat diversity, and positive correlation with overstory vegetation (*P* < 0.001, *P* < 0.001, *P* = 0.043, respectively; Table 2).

**Table 2.** Results of the AICc based multimodel inference of mean altitudinal range.

| Parameter | Model-averaged estimates | SE | P-value | Importance value |
| --- | --- | --- | --- | --- |
| **Mean Altitude Range** |  |  |  |  |
| Intercept | 1330.99 | 42.40 | < **0.001** *** | - |
| Minimum temperature | -24.05 | 5.06 | < **0.001** *** | 1.00 |
| Habitat diversity | -65.79 | 15.88 | < **0.001** *** | 1.00 |
| Understory vegetation | 3.33 | 7.47 | 0.658 | <0.001 |
| Midstory vegetation | -0.39 | 7.82 | 0.961 | <0.001 |
| Overstory vegetation | 3.90 | 6.8 | **0.043** * | <0.001 |

Candidate models included those with Akaike weights within 10% of the highest value. \*\*\* $P < 0.001$ ;
\*\* $P < 0.01$ ; \* $P < 0.05$ . SE = standard error.

### Testing of the non-directional rescue effect

We demonstrated the non-directional rescue effect using the intensity of species inflow. We found that species richness showed a tendency of increasing with increasing of distance between mean mid-points (Fig 2). A quadratic curve represented that the intensity of species inflow increased up to 300 m; however, it slightly decreased after 300 m (R^2^ = 0.19, *P* < 0.001; Fig 2.b). An exponential curve showed a same value of R^2^ with a quadratic curve. The lowest value of R^2^ was a linear curve (R^2^ = 0.16, *P* < 0.001; Fig 2.a).

**Fig 2.**
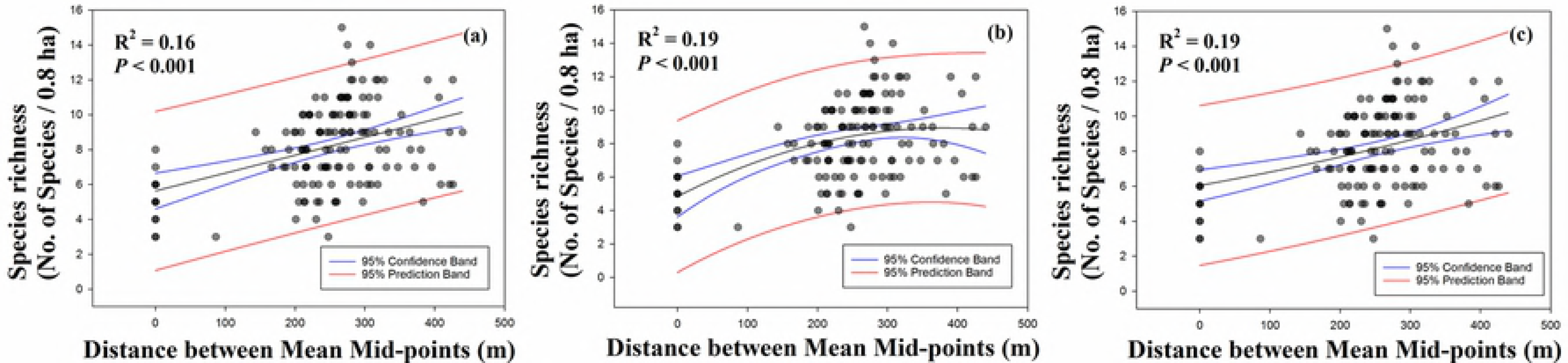
Relationship between species richness and distance between mean midpoints. **(a)** linear, (b) quadratic, and (c) exponential curves were represented.

### Effect of environmental factors on the range-size distribution group

We utilized more detailed methodology to identify which range-size distribution group increased species richness. From analysis of independent samples t-test, we found that the 2nd and 3rd quartile species contributed to increasing species richness (Fig 3). The 2nd and 3rd quartile species showed a significant differences in species richness between present and absent of each range-size quartile species (*P* = 0.002 and *P* = 0.009, respectively; Fig 3). Whereas, the 1st and 4th quartile species did not show a significant differences in species richness between present and absent (*P* = 0.447 and *P* = 0.195, respectively; Fig 3). The 4th quartile species showed substantial difference in the value of mean between present and absent (mean = 8.196 ± 2.443 and mean = 5.000 ± 0, respectively; Fig 3); however, the 4th quartile species did not show a significant difference because the most of areas were contributed by the 4th quartile species (present, n = 141 and absent, n = 1).

**Figure 3.**
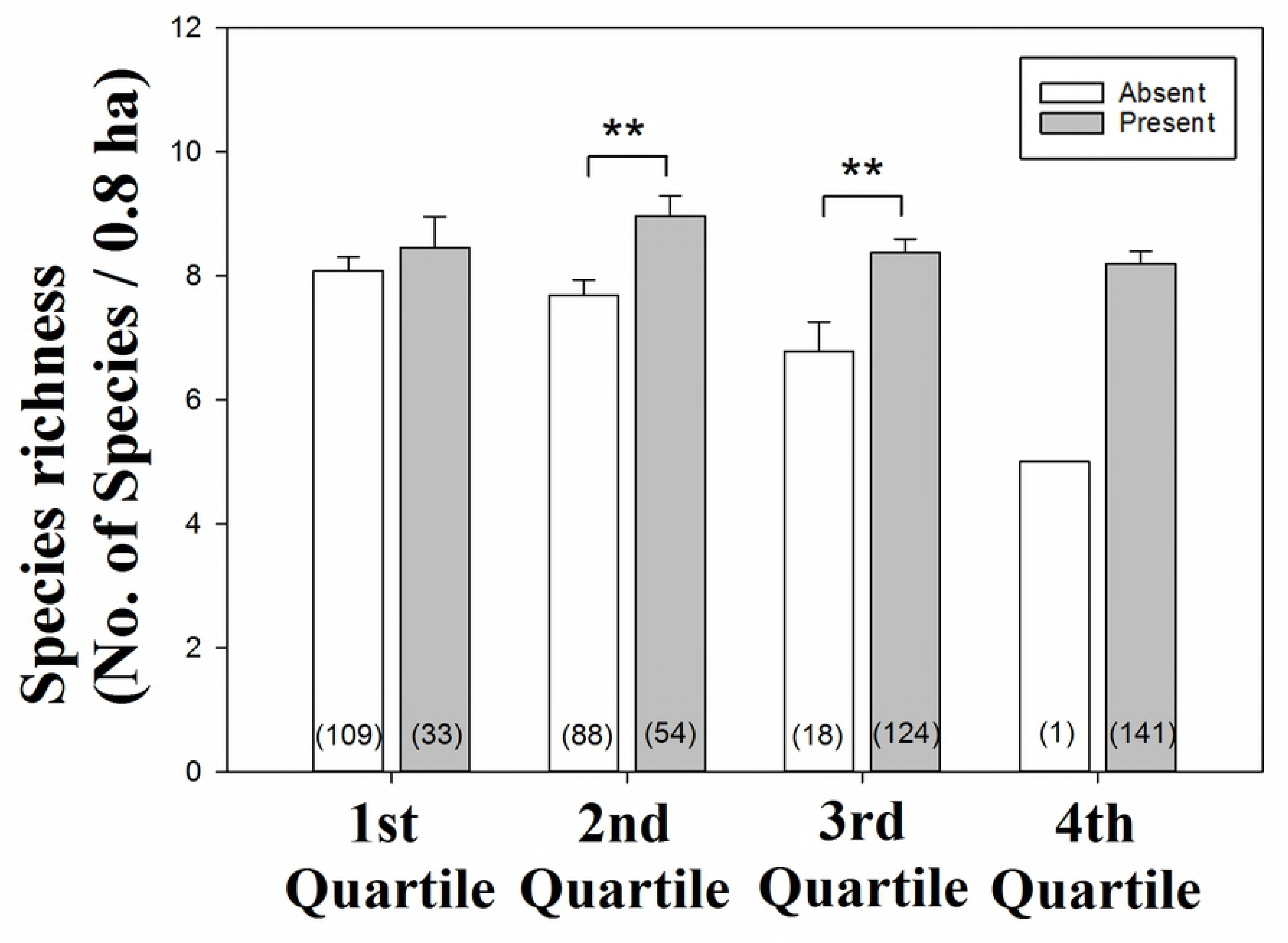
Effect of present and absent of each range-size quartile species on species richness. Vertical lines on the bars indicate SE (standard error) and bracket indicates sample size. \*\*\**P* < 0.001; ** *P* < 0.01; * *P* < 0.05.

To identify the effect of environment factors on 2nd quartile species richness, we utilized model selection and multimodel inference. A set of candidate models with seven combinations of five variables represented two supported models (Table 3). The best model included only the minimum temperature (*w*_i_ = 0.609; Table 3). Adding habitat diversity to the best model led to a 1.8-fold decrease in Akaike weight (*w*_i_ = 0.609 vs *w*_i_ = 0.331; Table 3). Adding vertical coverage of vegetation to the best model led to a 14.5-fold decrease in Akaike weight (*w*_i_ = 0.609 vs *w*_i_ = 0.042; Table 3). From the multimodel inference results, 2nd quartile species richness was influenced only by the minimum temperature (*P* < 0.001; Table 4)

**Table 3.** Model selection results for predicting 2nd quartile species richness based on minimum temperature, vertical coverage of vegetation, and habitat diversity.

| Response variables | Candidate models | AICc | ΔAICc | df | $w_i$ |
| --- | --- | --- | --- | --- | --- |
| Species richness of 2nd quartile | (Best model) Intercept + MT | 275.2 | 0 | 3 | 0.609 |
|  | Intercept + MT + HD | 276.5 | 1.2 | 4 | 0.331 |
|  | Intercept + MT + OV + MV + UV | 280.6 | 5.3 | 6 | 0.042 |
|  | (Full model) Intercept + MT + OV + MV + UV + HD | 282.2 | 7.0 | 7 | 0.018 |
|  | Intercept + HD | 298.0 | 22.8 | 3 | <0.001 |
|  | Intercept + OV + MV + UV + HD | 302.2 | 27.0 | 6 | <0.001 |
|  | Intercept + OV + MV + UV | 321.2 | 46.0 | 5 | <0.001 |

Coverage of understory vegetation = UV, midstory vegetation = MV, overstory vegetation = OV, minimum temperature = MT, habitat diversity = HD

**Table 4.** Results of the AICc-based multimodel inference of 2nd quartile species richness.

| Parameter | Model-averaged | SE | P-value | Importance |
| --- | --- | --- | --- | --- |

|  | estimates |  |  | value |
| --- | --- | --- | --- | --- |
| **Species richness of 2nd quartile** |  |  |  |  |
| Intercept | -1.759 | 0.35 | <0.001*** | - |
| Minimum temperature | 0.241 | 0.04 | <0.001*** | 1.00 |
| Habitat diversity | 0.131 | 0.14 | 0.352 | 0.35 |

Candidate models included those with Akaike weight within 10% of the highest value. \*\*\* $P < 0.001$ ;
\*\* $P < 0.01$ ; \* $P < 0.05$ . SE = standard error.

We identified two supported models that showed Akaike weights within 10% of the highest value (Table 5). The best model included habitat diversity and vertical coverage of vegetation (*w*_i_ = 0.625; Table 5). The second ranked model was the full model that also contained the minimum temperature (Table 5). The model containing vertical coverage of vegetation was 16.9 times more likely to be the best explanation for the 3rd quartile species richness (*w*_i_ = 0.625 vs *w*_i_ = 0.037; Table 5). When habitat diversity was included in the 3rd quartile species richness model, Akaike weights were 8.5 times higher than those eliminated in the model (*w*_i_ = 0.289 vs *w*_i_ = 0.034; Table 5). Multimodel averaged parameter estimates including the two supported models in the 3rd quartile species richness represented a positive relationship with habitat diversity, and understory and overstory vegetation (*P* = 0.001, *P* = 0.005, and *P* = 0.032, respectively; Table 6).

**Table 5.**
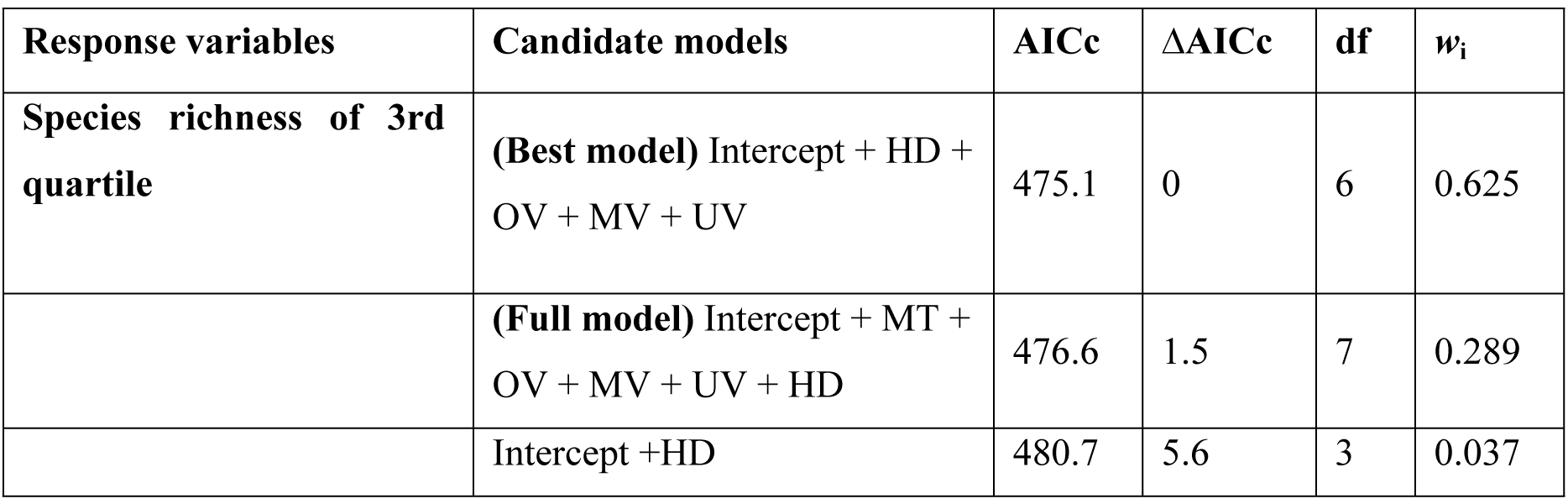
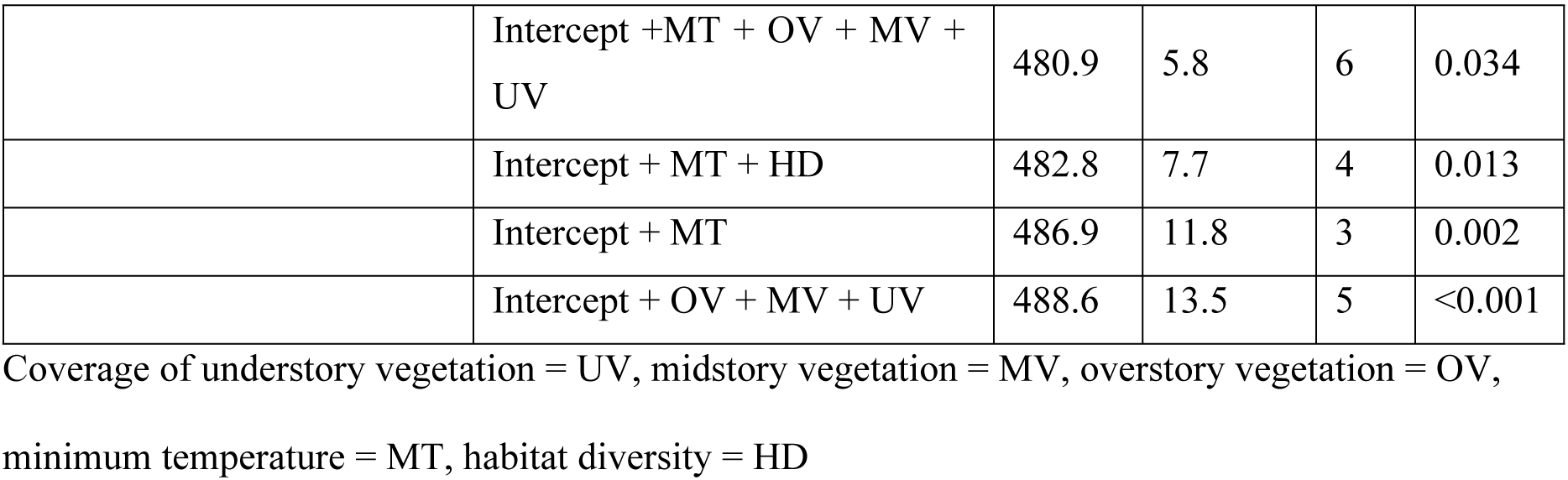
Model selection results for predicting 3rd quartile species richness based on minimum temperature, vertical coverage of vegetation, and habitat diversity.

**Table 6.**
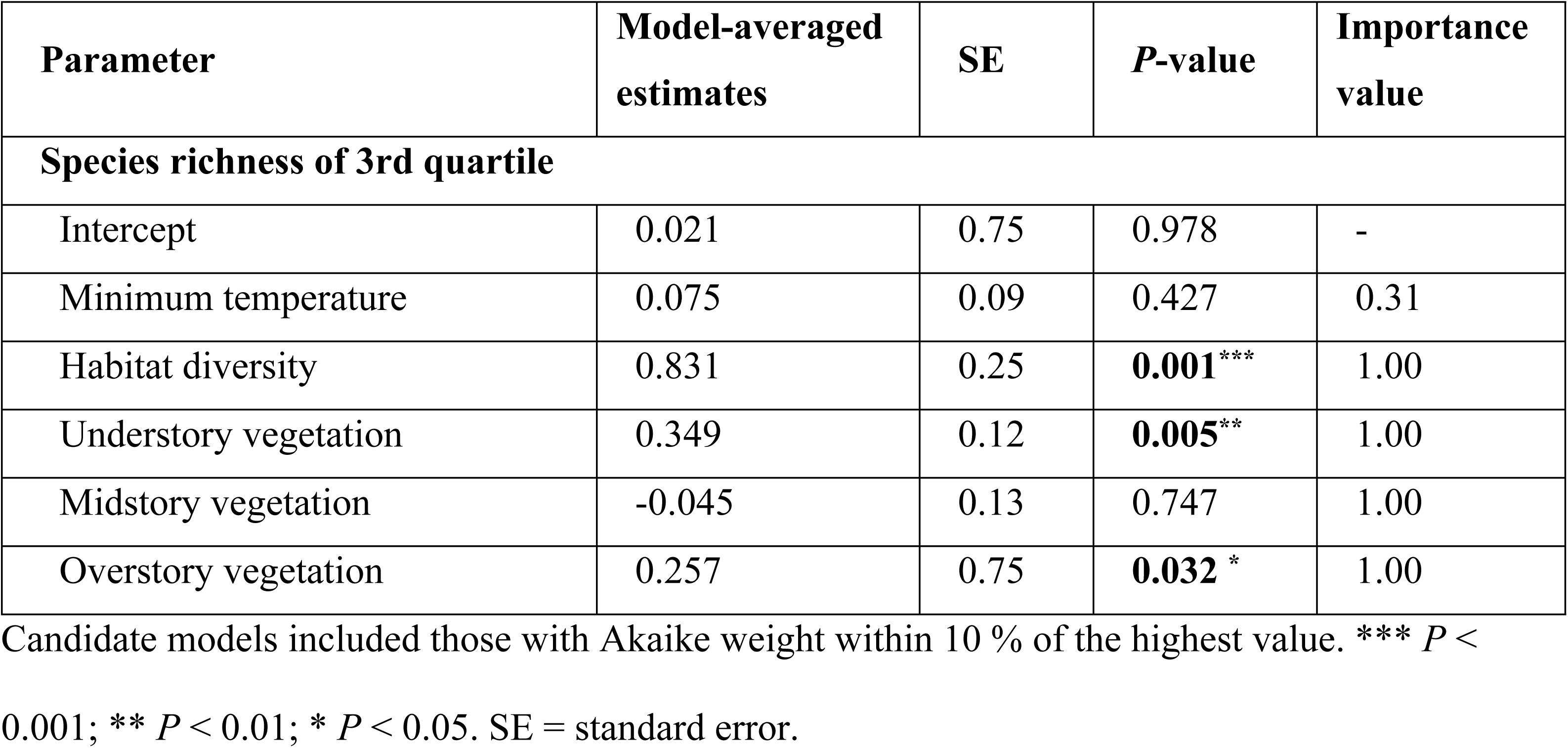
Results of the AICc-based multimodel inference of 3rd quartile species richness.

| Parameter | Model-averaged estimates | SE | P-value | Importance value |
| --- | --- | --- | --- | --- |
| **Species richness of 3rd quartile** |  |  |  |  |
| Intercept | 0.021 | 0.75 | 0.978 | - |
| Minimum temperature | 0.075 | 0.09 | 0.427 | 0.31 |
| Habitat diversity | 0.831 | 0.25 | **0.001**<sup>***</sup> | 1.00 |
| Understory vegetation | 0.349 | 0.12 | **0.005**<sup>**</sup> | 1.00 |
| Midstory vegetation | -0.045 | 0.13 | 0.747 | 1.00 |
| Overstory vegetation | 0.257 | 0.75 | **0.032**<sup>*</sup> | 1.00 |

Candidate models included those with Akaike weight within 10 % of the highest value. \*\*\* $P < 0.001$ ; \*\* $P < 0.01$ ; \* $P < 0.05$ . SE = standard error.

## Discussion

### Species range-size distribution (Altitudinal Rapoport’s rule)

McCain and Bracy Knight [24] asserted that the altitudinal Rapoport’s rule is not pervasive in vertebrate taxa. In bird taxa, approximately 45% strong support for the altitudinal Rapoport’s rule was found, whereas no trend had approximately 55% support [24]. The reason why the altitudinal Rapoport’s rule is not supported is owing to differences in sampling effort, habitat type, methodology, geographical scale, and mechanism used [7–10,24,25]. Stevens [6] stated that compared to sampled point studies, regional surveys are more likely to be biased owing to unequal sampling. If an intensive survey is undertaken only in one elevation band, then species richness will be biased upward and altitudinal range will be biased downward [6]. Thus, in the present study, we conducted a field survey that utilized identical sampling intensity (S2 Table) and sampled using a point count survey method. Additionally, we performed the field survey restrictively in a mixed or deciduous forested area [17]. Food sources are the most influential factor on the distribution of birds during the breeding season. Conifer forests can have a completely different species composition; thus, we minimized the effect of any differences caused by vegetation type by confining the study to mixed or deciduous forested areas. McCain and Bracy Knight [24] stated that mountains above 23° N latitude detected significantly stronger support for the altitudinal Rapoport’s rule. Our study site was located at above 23° N latitude (S2 Table), thus the above-mentioned geographical features were influential. Although the present study was conducted in a relatively small mountainous area having a low elevation range, our results are similar to those found for the altitudinal Rapoport’s rule.

### Climatic and ambient energy hypotheses

To understand the phenomenon that higher altitudinal species have wider altitudinal ranges, we tested the underlying mechanisms, i.e., the climatic and ambient energy hypotheses. According to the climatic hypothesis, we assumed that species that have a broader physiological thermal tolerance also have a wider altitudinal range-size distribution [3,6,11], thus altitudinal range-size distribution should show a negative relationship with minimum temperature. Our results showed that minimum temperature was the most important factor among the variables and that altitudinal range-size distribution increased in regions with severe physiological thermal tolerance (Table 1 and 2), thus supporting the climatic hypothesis.

According to the ambient energy hypothesis, greater habitat heterogeneity and primary productivity have negative relationships with range-size distributions [15]. To understand this relationship, we preliminarily needed to understand the role of species richness. In general, greater habitat heterogeneity and primary productivity could promote higher numbers of species [15,26]. As stated by Rapoport’s rule, in the equatorial and low altitudinal regions there is higher species richness owing to greater habitat heterogeneity and primary productivity without interspecific interactions [6,15]. Limited food resources and habitat competition lead to the determination of species range-size distributions. Thus, based on the mechanisms of Rapoport’s rule and the ambient energy hypothesis, our results showed a negative relationship between range-size distribution and habitat diversity (habitat heterogeneity). However, our results also showed a positive relationship with the coverage of overstory vegetation (primary productivity) (Table 2), thus the results did not support the ambient energy hypothesis. Here, we found some logical error. Rapoport’s rule describes range-size distribution based on the assumption that species richness is higher in equatorial or low altitudinal regions [6]. In the present study, the range-size distribution showed an increasing pattern with increasing altitude according to the altitudinal Rapoport’s rule (Fig 1); however, species richness showed a mid-peak pattern [17]. Therefore, these two results did not show proper logical flow. To grasp the relationships among the patterns of species richness, range-size and environmental factors more closely, we must understand the rescue effect and the effect of environmental factors on quartile species. Our results could explain the pattern of range-size distribution from the effect of climatic and habitat diversity; however, the relationship between species richness and range-size distribution could not be explained.

### Non-directional rescue effect

Almeida-Neto et al. [16] argued that Steven’s model predicted a peak in species richness at lower elevations owing to asymmetric inflation in species richness from higher to lower elevations. The non-directional model is expected to generate a peak at mid-elevations based on the mid-domain effect [16,27–29]. A non-directional rescue effect proposes the inflow of species from both directions (high and low elevations). Thus, we identified the intensity of species inflow using this new method. From testing of the non-directional rescue effect, the species richness showed a tendency of increasing with increasing of species inflow (Fig 2). According to our prediction, the reason for higher species richness at mid-elevation was owing to species inflow from other areas apart from the mid-altitude area. Thus, our results supported the non-directional rescue effect. To demonstrate the reason behind higher species richness at mid-elevation in relation to species range-size distribution, we used the quartile method.

### Effect of environmental factors on the range-size distribution group

We found that the 2nd and 3rd quartile species contributed to the increased species richness at mid-elevation (Fig 3). From the results of distribution patterns of each quartile species, we found that the distributions of the 4th quartile species were skewed toward high altitudes, the 2nd quartile species were skewed toward low altitudes, and the 3rd and 1st quartile species were distributed over the entire altitudinal range (Fig 4). As shown in Fig 4, the 4th quartile species tended to be skewed toward high altitudes but were distributed over a wide altitudinal range similar to generalist species [30], thus the 4th quartile species did not affect species richness compared to the other quartile species. Because the most of areas were equally contributed by the 4th quartile (Fig 3 and 4). The 1st quartile species did not contribute to species richness either, because these species showed only a small number of detections and possessed specialist species features (Fig 3) [30,31]. Thus, the distributions of the 2nd and 3rd quartile species contributed to the increase of species richness. The cause of increasing range-size distribution was attributed to the distribution patterns of the quartile species, and the 2nd and 3rd quartile species were the main cause of the increasing species richness and the non-directional rescue effect.

**Fig 4.**
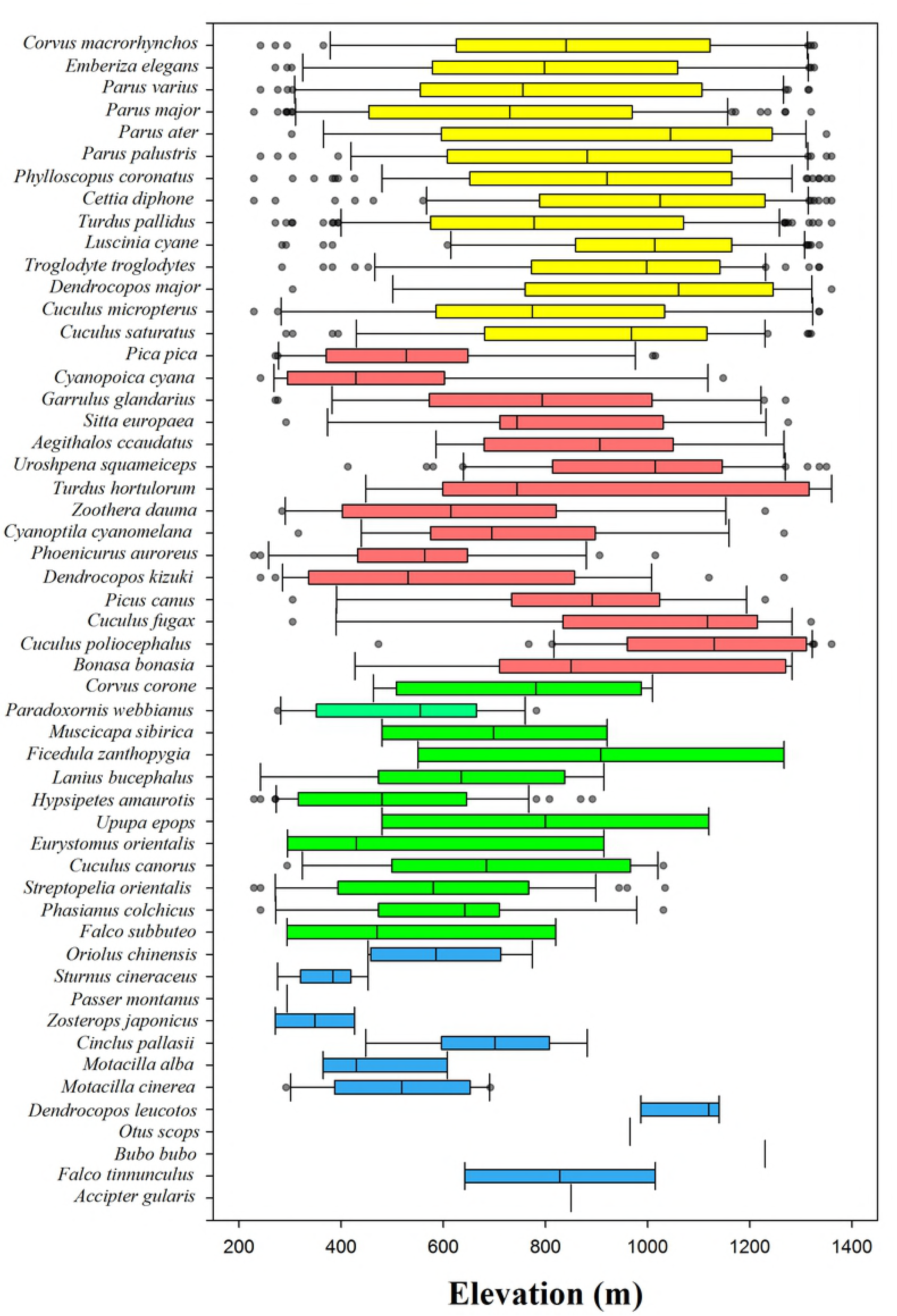
Species distribution of each quartile along altitudinal gradient. Different colors indicate each quartile group. Blue = 1st quartile species, green = 2nd quartile species, red = 3rd quartile species, yellow = 4th quartile species.

We identified the effect of environmental factors on the 2nd and 3rd quartile species richness that contributed to the mid-peak of species richness. In general, the 1st and 2nd quartile species are composed of species having a narrow altitudinal range-size distribution. Thus, we assumed that the 1st and 2nd quartile species have a narrow physiological thermal tolerance, and possess thermal specialist species features [32,33]. However, we found that the only 2nd quartile species preferred a range of warm temperatures (Table 4; Fig 4), and possessed thermal specialist species features. Whereas, the 1st quartile species was influenced by only habitat diversity and distributed over the entire altitudinal range (S3 Table; Fig 4), was identified having a features of habitat specialist species. Meanwhile, the 3rd and 4th quartile species are composed of species having a wide altitudinal range-size. Thus, we assumed that the 3rd and 4th quartile species are not influenced by habitat and temperature, and possess generalist species features [34,35]. However, we found that the 3rd quartile species was influenced by habitat heterogeneity and primary productivity. Whereas, the 4th quartile species was influenced by primary productivity and minimum temperature (Table 6; S3 Table). A previous study conducted on latitudinal differences, the 3rd and 4th quartile species were strongly influenced by primary productivity compared to other quartile groups [2], showed a coincidence with our results. To achieve a better understanding of these patterns, competition among species related to niche are required. From these results, we determined that the cause of mid-peak pattern of species richness was not inflow of habitat specialist species [14,15], but owing to the influence of minimum temperature, habitat heterogeneity, and primary productivity on the distribution of the 2nd and 3rd quartile species.

## Conclusions

Altitudinal range-size distribution increased with increasing altitude and showed a negative relationship with minimum temperature and habitat diversity and a positive relationship with coverage of vegetation. These results support the altitudinal Rapoport’s rule and climatic hypothesis; however, they do not fully support the ambient energy hypothesis. There was some logical error between the Rapoport’s rule and mid-peak pattern of species richness. Thus, we tested the non-directional rescue effect, and the results supported this effect. Using the quartile method, we found that the 2nd and 3rd quartile species richness were the main contributors to the mid-peak of species richness and the non-directional rescue effect. The 2nd quartile species richness was influenced by minimum temperature and possess thermal specialist species features, and the 3rd quartile species was influenced by habitat heterogeneity and primary productivity. Although altitudinal range-size distribution results were similar to the altitudinal Rapoport’s rule, the mid-peak pattern of species richness could not be explained by the underlying mechanism of altitudinal Rapoport’s rule. However, the non-directional rescue effect could explain a mid-peak pattern of species richness along altitudinal gradient. The reason why the non-directional rescue effect and mid-peak of species richness was because of the influence of proper temperature and habitat complexity on the distribution of the 2nd and 3rd quartile species.

## Supporting Information

**S1 Table. Pearson’s correlations between climatic (maximum temperature and minimum temperature), vertical (coverage of understory, midstory, and overstory vegetation) and horizontal habitat heterogeneity (habitat diversity).** Bold = correlated predictor (r ≥ |0.7|).

**S2 Table. Information of the study plots along altitudinal gradient.**

**S3 Table. Results of the AICc based multimodel inference of species richness of the 1st and 4th quartiles.**

